# Conformation-specific Design: a New Benchmark and Algorithm with Application to Engineer a Constitutively Active Map Kinase

**DOI:** 10.1101/2025.04.23.650138

**Authors:** Jacob A. Stern, Siba Alharbi, Anandsukeerthi Sandholu, Stefan T. Arold, Dennis Della Corte

## Abstract

A general method for designing proteins with high conformational specificity is desirable for a variety of applications, including enzyme design and drug target redesign. To assess the ability of algorithms to design for conformational specificity, we introduce MotifDiv, a benchmark dataset of 200 conformational specificity design challenges. We also introduce CSDesign, an algorithm for designing proteins with high preference for a target conformation over an alternate conformation. On the MotifDiv benchmark, CSDesign designs protein sequences that are predicted to prefer the target conformation. We apply this method *in vitro* to redesign human MAP kinase ERK2, an enzyme with active and inactive conformations. Out of two designs for the active conformation, one increased activity sufficiently to retain activity in the absence of activating phosphorylations, a property not present in the wild type protein.

## 1 Introduction

### 1.1 Inverse folding

Inverse folding (protein sequence design for a given structure) has advanced rapidly with both physics-based and deep learning methods. Traditional approaches like Rosetta fix a backbone and search for low-energy sequences using rotamer packing algorithms (Alford et al., 2017). More recently, deep learning methods have achieved remarkable performance in inverse folding. ProteinMPNN is a graph-neural-network model that generates sequences conditioned on the 3D coordinates of a protein backbone. It significantly outperforms Rosetta in sequence recovery (52% vs 33% on native structures) and designs sequences that fold experimentally. Another approach uses AlphaFold2’s structure prediction network for design. By inverting AlphaFold – e.g. via gradient descent or MCMC on input sequences – researchers can optimize sequences to fold into a target structure (Goverde et al., 2023).

### 1.2 Conformational specificity as a design objective

Designing a protein not just to fold stably, but to prefer one specific conformation or functional state over alternatives, is a key goal in many applications:

#### Enzyme design and constitutive activity

Many enzymes have active and inactive conformations (e.g., due to regulatory domains or flexible loops). Designing a constitutively active enzyme often means stabilizing the active state so it no longer requires its natural trigger. For instance, Dowling et al. (2023) computationally designed mutants of cyclic GMP-AMP synthase (cGAS) that adopt the active conformation without DNA binding. Using a two-state design strategy, they biased the sequence energy landscape toward the active state and away from the inactive form. This illustrates how multi-state design can stabilize one conformation (active) at the expense of another (inactive) to achieve continuous activity.

#### Drug targeting and allosteric states

Many drug targets (ion channels, kinases, GPCRs) undergo conformational changes between “open” and “closed” or active/inactive states. Protein variants that maintain one conformation with high specificity make it possible to screen and identify drug hits against a specific conformation, making it possible to tactically target specific protein functions.

#### Conformational specificity as an ML design objective

Conformational specificity is increasingly a consideration in machine learning-based protein design. This commonly takes the form of post-hoc filtering generated sequences. For example, a design framework might use AlphaFold2 as a referee: for a candidate sequence design, if AlphaFold confidently predicts the target structure and not alternative folds, the sequence is kept. This approach was successful for ProteinMPNN; sequences that did so were far more likely to fold experimentally Goverde et al. (2024). Another ML approach, which we investigate here, is direct generative modeling of sequences specific to one conformation over another.

### 1.3 This work

The contributions of this work are as follows:

- We introduce MotifDiv, a dataset of 200 conformational specificity design challenges within the Protein Data Bank (PDB).
- We introduce CSDesign, an inference-time adaptation of ProteinMPNN for designing proteins with conformational specificity.
- We show that CSDesign successfully designs proteins to prefer target conformations on the MotifDiv dataset.
- We use CSDesign to redesign human ERK2 kinase to prefer its active conformation, and successfully convert the natively-inactive wild type into a constitutively active enzyme.

## 2 Methods

### 2.1 Boltzmann conformational specificity objective

We adopt the Boltzmann-motivated probabilistic definition of conformational specificity introduced by (Stern et al., 2023), which quantifies the preference of a sequence for a target conformation relative to an alternate folded state:

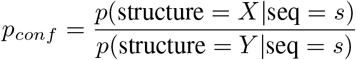

When optimizing this objective, applying Bayes’ Theorem simplifies the objective to a tractable probability ratio for an inverse folding model (see section A.1.1):

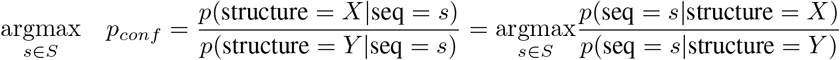

We employ ProteinMPNN Dauparas et al. (2022) as a model for *p*(seq = *s*|structure = *X*), using a decoding algorithm detailed in section A.1.2.

We study the effect of inverse folding with a conformational specificity objective 1) *in silico* on the MotifDiv benchmark and 2) *in vitro* for a human Extracellular Signal-Regulated Kinase 2 (ERK2), which changes conformation upon phosphate binding.

### 2.2 Evaluation of designs *in silico*

To evaluate the ability of the model to design proteins with high conformational specificity, we filtered the PDB to create a subset of design challenges. The MotifDiv dataset is a selection of PDB pairs with high structural homology except for significant divergence within a 10-residue motif. The creation of the MotifDiv dataset is detailed in section A.2. The result is 200 single-chain domain pairs (400 total domains).

In *in silico* studies, we redesign the sequence of each instance within each pair for a total of 400 designs per tested model. We then use ESMFold (Lin et al., 2023) to predict the structure of each designed sequence and compute the scaffold-aligned motif RMSD between the predicted and the target structure, as well as between the predicted and the alternate structure. The results are shown in Table 1 and Figure 3.

**Table 1:** The first two metrics assess the ability of the models to design with a preference for the target conformation. The second two metrics assess the ability of one model to outperform the other. Experiments are described in detail in section 3.1.

| Statistic | Effect Size (Å) | t-statistic | p-value |
| --- | --- | --- | --- |
| $\text{RMSD}_{diff\_csd}$ | 2.01 | 6.42 | $3.8\text{e-}10$ |
| $\text{RMSD}_{diff\_mpnn}$ | 1.77 | 5.10 | $1.0\text{e-}07$ |
| $\text{RMSD}_{diff\_comp}$ | 0.245 | 1.35 | 0.17 |
| $\text{RMSD}_{sum\_comp}$ | 0.503 | 1.48 | 0.14 |

### 2.3 Evaluation of designs *in vitro*

MAP kinases require a dual phosphorylation on a tyrosine and serine/threonine in their ‘activation loop’. This phosphorylation is carried out by an upstream kinase (MAP kinase kinase; MAPKK). The attached phosphate groups engage intramolecular interactions with the core of the kinase, resulting in a specific stable structuring of the activation loop and overall active kinase conformation. The activation loop then can serve as a basis for substrates.

We design 4 variants of human MAP kinase ERK2: 2 variants preferring the active conformation (PDB 2ERK) (Canagarajah et al., 1997) and 2 variants preferring the inactive conformation (PDB 4GSB). Residues 169-186 exhibit greatest structural variation between the two conformations, so we redesign residues in this region. For each conformation we use two strategies - 1) redesigning only residues in a linear region consisting of residues 169-186, and 2) redesigning all residues which fall within an 8Å spatial radius of these residues.

Protein designs are commercially synthesized, recombinantly expressed in E. coli, and purified using standard procedures (see A.3.2). Catalytic activity of the purified proteins is assessed in vitro using the ADP-Glo kinase assay.

## 3 Results

### 3.1 *In silico* Evaluation on the MotifDiv dataset

We found that both CSDesign and ProteinMPNN preferred their target conformations with statistical significance (see Figure 1). We performed a one-sample t-test on the statistic *RMSD*_*diff*_ = *RMSD*(*pred, pro*) −*RMSD*(*pred, alt*). CSDesign averaged 2.30 Å, with a p-value of 1.99e-11, showing preference for the target conformation across the dataset. Similarly, ProteinMPNN averaged 1.77 Å with a p-value of 1.0e-7, likewise succeeding in designing to the target conformation.

**Figure 1.**
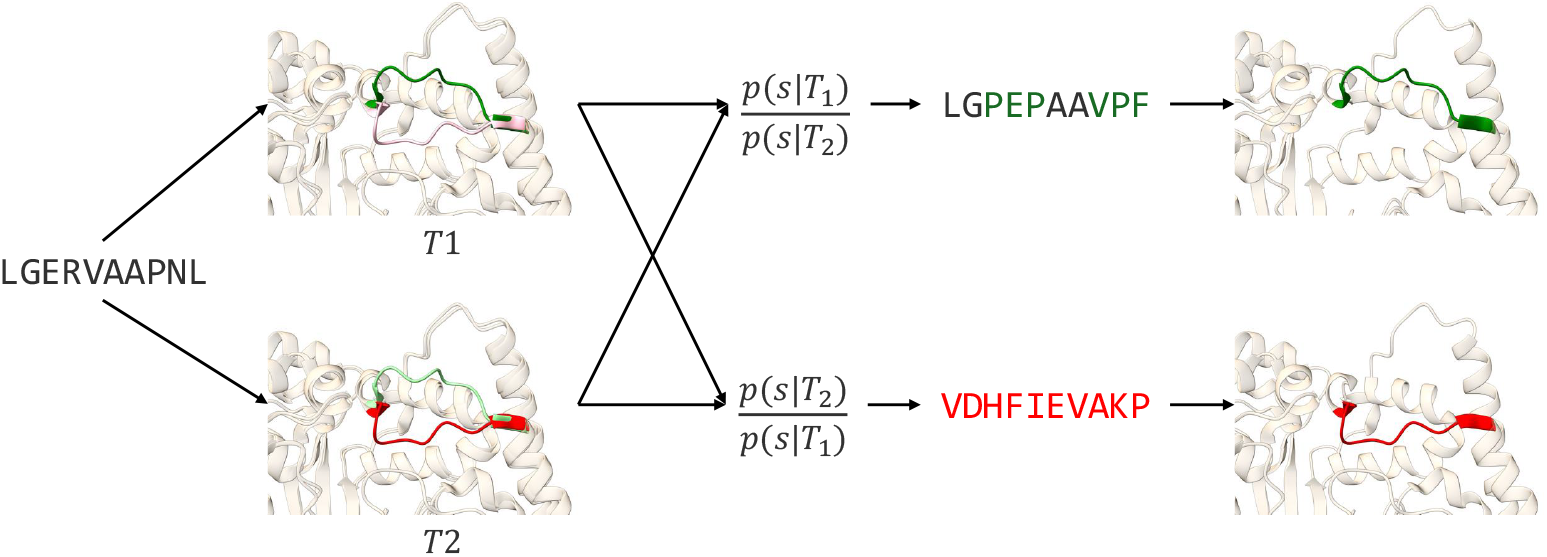
CSDesign is an inference-time sequence decoding algorithm for designing protein sequences with strong conformational preference. Given two reference structures, the model uses the ratio of sequence probabilities to design sequences that prefer one conformation or the other.

**Figure 2.**
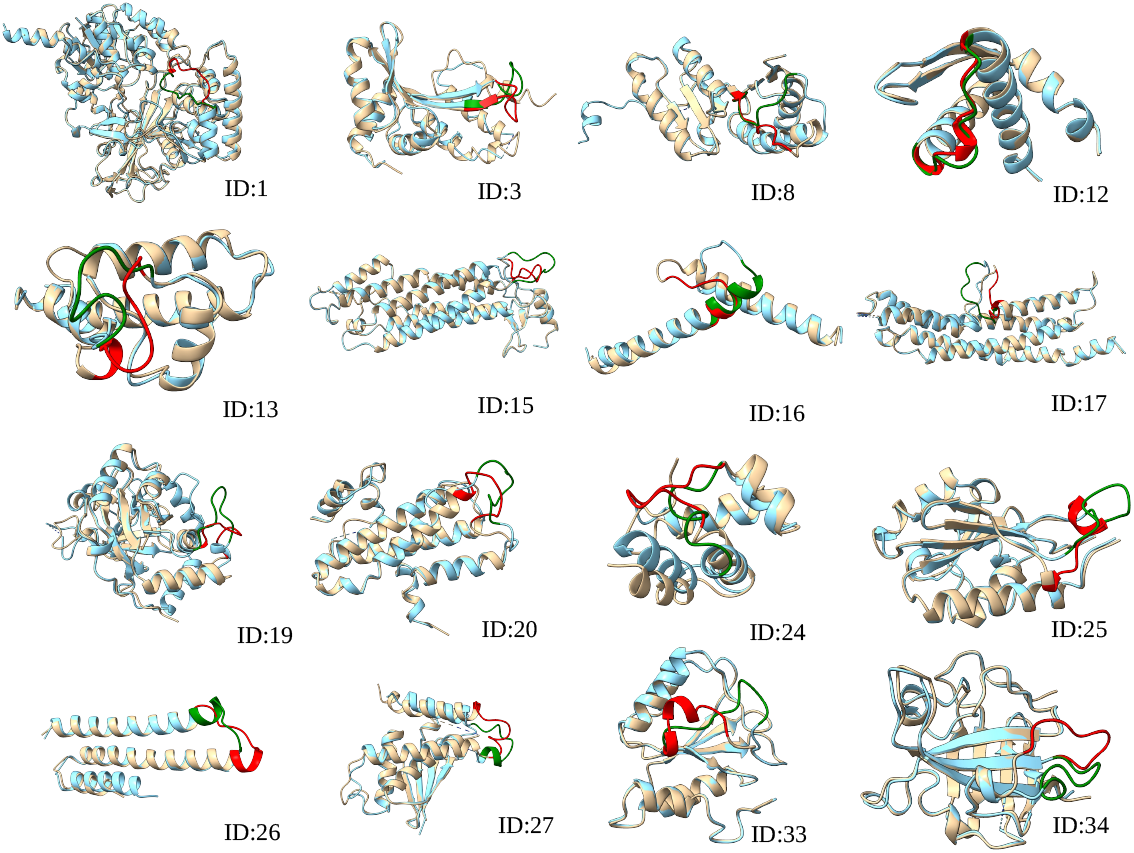
Selected examples from the MotifDiv dataset. Matches were selected such that they contained high structural homology in the scaffold region (brown and blue), but conformational divergence within a small motif region (green and red).

**Figure 3.**
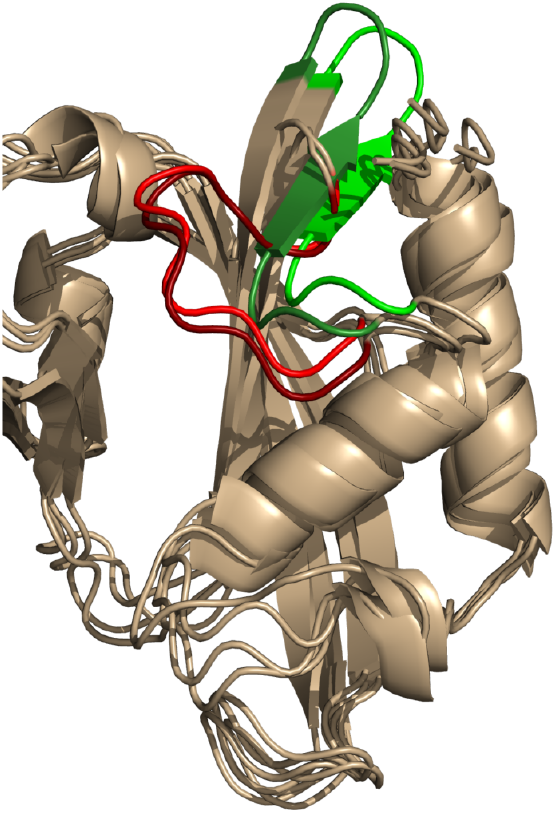
Figure showing the MotifDiv pair with lowest summed-motif RMSD for CSDesign. Dark green represents one target motif, dark red represents the other target motif. Light green and light red represent the predicted structures of the sequences designed to prefer these motifs, respectively.

Next, we compared CSDesign to ProteinMPNN with a one-sample t-test on the statistic *RMSD*_*diff*_ *_*_*comp*_ = *RMSD*_*diff mpnn*_ −*RMSD*_*diff*_ *_*_*csd*_ to evaluate whether the improvement in specificity of CSDesign over ProteinMPNN was statistically significant, and found that it was not.

Finally, we compared CSDesign to ProteinMPNN on paired conformers. We performed a one-sample t-test on the statistic *RMSD*_*sum*_*_*_*comp*_ = *RMSD*_*summed*_*_*_*mpnn*_ −*RMSD*_*summed*_*_*_*csd*_ where 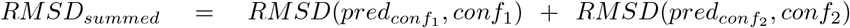 and 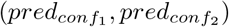 are separate predictions for each conformer in a pair (*conf*_1_, *conf*_2_). In this comparison, we again found no statistically meaningful improvement of CSDesign over ProteinMPNN.

### 3.2 In vitro Evaluation on ERK2 Redesign

Proteins CSD101 and CSD102 were designed to prefer the inactive conformation, corresponding to PDB 4GSB. CSD101 did not demonstrate measurable activity above a baseline in the ADP-GLO assay, and CSD102 was not expressed successfully. The wild type variant also had no measurable activity over the control, as expected.

Proteins CSD103 and CSD104 were designed to prefer the active conformation corresponding to PDB 2ERK. CSD103 did not demonstrate detectable activity; however, CSD104 illuminated, even in the absence of ERK2 phosphorylation (see Table 2). This signals success in designing an ERK2 variant that prefers the active conformation.

**Table 2:** *in vitro* screening of design variants in the ADP-GLO assay identifies a constitutively active ERK2 variant

| Sequence ID | CSDesign preferred conformation | Edit distance to WT100 | Expression | ADP-Glo % Activity w/o PO<sub>3</sub> |
| --- | --- | --- | --- | --- |
| Control | - | - | - | 1.65 ± 0.33 |
| WT100 | - | 0 | Yes | 1.48 ± 0.24 |
| CSD101 | Inactive | 20 | Yes | 2.23 ± 0.03 |
| CSD102 | Inactive | 55 | No | - |
| CSD103 | Active | 12 | Yes | 2.92 ± 0.34 |
| CSD104 | Active | 23 | Yes | 46.22 ± 9.03 |

*In silico* metrics also indicated that CSD101 and CSD102 adhered more to the inactive conformation and CSD103 and CSD104 adhered more closely to the active conformation, as shown in table S3 and figure S7.

## 4 Conclusion

This work introduces the first benchmarking dataset for conformation-oriented protein design, introduces a new algorithm for conformation-specific sequence design, and pioneers its use in engineering a constitutively active kinase.

In this work, we redesign the sequence of the same region in which we want to modulate the conformation. A promising frontier is “allosteric design”, in which the design region differs from the target conformation region. This could make it possible to alter the conformation of the target region without sacrificing the functional properties of its amino acid sequence. Allosterically related residues could be selected by a method like Kannan et al. (2024), which demonstrated an unsupervised method of identifying allosteric relationships from attention maps.

We also note that the MotifDiv dataset is not limited to assessing conformation-specific design. It could also be used to assess performance for a complementary problem, multi-state protein design Sauer et al. (2020).

This work lays the groundwork for future progress in conformation-specific protein design, with applications in drug discovery and enzyme engineering.

## Data and Code availability

Code for the CSDesign algorithm is available at https://github.com/dellacortelab/cs_design. The MotifDiv dataset is available at https://github.com/dellacortelab/motif_div. Experimental data is available from the corresponding author upon request.

## Acknowledgments

The authors would like to thank David Wingate for discussion and ideation for this work.

Experimental research was supported by the Bioscience Core Lab, and ACL Proteomics Core Lab at King Abdullah University of Science & Technology (KAUST) in Thuwal, Saudi Arabia. The work by STA, SA, and AS was supported by the King Abdullah University of Science and Technology (KAUST) through the baseline fund to STA and under Award No. FCC/1/5932-09-01. The work of DDC was supported by the National Institute of General Medical Sciences of the National Institutes of Health under award number R15GM155803.

## A APPENDIX

### A.1 CSDesign methods

#### A1.1 Objective derivation

Stern et al. (2023) introduces a Boltzmann-motivated definition of conformational specificity of a protein sequence *s*:

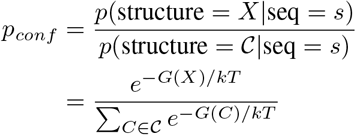

Where *X* is the macrostate corresponding to the folded conformation of interest and *C* is the macrostate subsuming all alternate folded conformations. If there is one known alternate conformation *Y* which dominates the denominator, then this definition simplifies to:

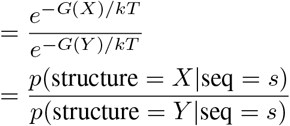

By applying Bayes’ rule to this objective and maximizing over sequence, the objective reduces to a probability ratio that is tractable for an unmodified inverse folding model:

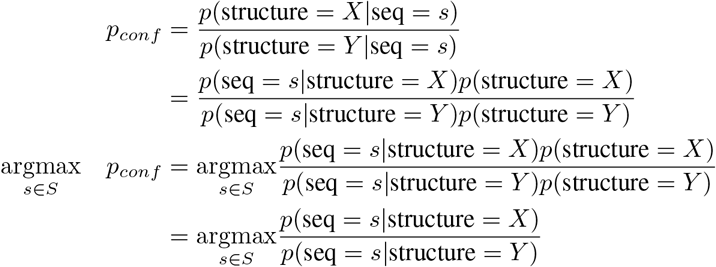

This objective is similar to Stern et al. (2023)^1^, but requires only one model, an inverse folding model of the form *p*(seq|structure = *X*) ^2^. We use ProteinMPNN (Dauparas et al., 2022), and the decoding algorithm is described in SI section A.2.1. We also observe that this algorithm has the same limitation described in Stern et al. (2023) -namely, that it relies on the argmax operator and thus requires some form of a greedy decoding scheme, making it unsuitable for sampling schemes commonly used in autoregressive models.

This derivation can similarly be applied to the motif/scaffold case, as is used in this paper:

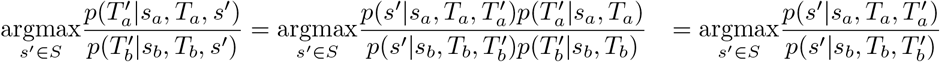

where subscripts *a* and *b* refer to conformations *a* and *b*, 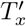 refers to the motif structure, *s*^*′*^ refers to the designed motif sequence, and *T*_*x*_ and *s*_*x*_ refer to the fixed scaffold structure and sequence.

#### A.1.2 Decoding

We can factorize the joint probability of a sequence as the product of conditional probabilities using the chain rule of probability. This reduces to a ratio of probabilities for each position in the sequence, where at each position the probability ratio gives a score for each amino acid, written as follows:

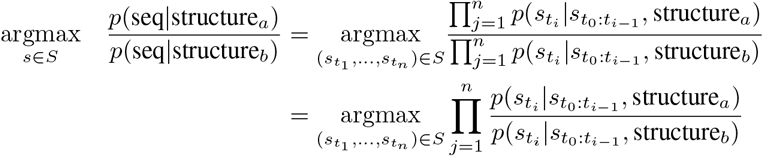

where {*t*_*i*_} is a decoding order and is a choice of the user. We can also hold any position in the sequence fixed, where *s*_*i*_ is assigned, *p*(*s*_*i*_) passes out of the argmax operator, and the subsequent tokens are conditioned on *s*_*i*_

### A.2 MotifDiv dataset methods

The dataset was generated by filtering domains from the PDB. The general objective was to identify pairs of structures with high structural homology in a “scaffold” region and high structural diversity in a 10-residue “motif” region. We used the CATH protein domain classification clusters Sillitoe et al. (2021) to narrow our search for matches within similar clusters.

Selection process 1:

- For each domain, we randomly selected 20 other domains from its CATH cluster.
- For each comparison domain, we retained it if it had over 90% full-sequence identity and 100% identity within at least one 10-residue region.
- We then stored the 10-residue, 100% sequence identity region with highest motif-aligned motif RMSD.
- We discarded all matches for that domain except the match with the highest motif RMSD.

We further filtered this selection to remove matches where:

- The total length was *>*650 residues.
- The motif of protein A sterically clashed with the scaffold of protein B or vice versa (under a scaffold alignment).
- The motif region was within 15 sequence positions of the N- or C-terminus.
- The motif region was within 10 sequence positions of a missing (disordered) residue.
- There were fewer than 20 scaffold residues.
- The domains were from the same protein.

We then de-duplicated on PDB id such that no PDB id occurs more than once in the dataset.

Finally, we sorted on motif RMSD and selected the 200 pairs with largest motif RMSD.

### A.3 ERK2 Study

#### A.3.1 Methods extended

##### Protein cloning, expression and purification

All four constructs were cloned by TWIST bioscience Ltd. in a pJEx411c vector with kanamycin antibiotic resistance. These plasmids were then transformed into E. coli BL21(DE3) competent cells and grown at 37°C in LB medium containing 50 *µ*g/ml kanamycin until the cell density reached an absorbance at 600 nm of 0.6 to 0.8, protein expression was induced with 0.25 mM IPTG for 16 h at 18°C. Cells were then harvested, centrifuged, and the cell pellet was resuspended in lysis buffer (50 mM Tris HCl pH 8.0, 500 mM NaCl, 10 mM imidazole, 2 mM Bme), 0.1% triton, a tablet of protease inhibitor, and benzonase. Cell suspensions were lysed using a sonicator on icy water bath and then centrifuged at 89,000 g for 30 mins to remove cell debris. The protein was purified from supernatant using a 5 ml HisTrap column (GE Healthcare). The proteins were eluted using 500 mM imidazole. Then all the proteins were passed through size exclusion chromatography column, Superdex 200 Increase 10/300 (GE Healthcare), equilibrated with the buffer containing 20 mM HEPES, pH 7.5, 200 mM NaCl, and 2 mM DTT. Proteins were concentrated using ultrafiltration membrane (Merck Millipore) with 30 kD MW cut-off for experiments and stored at −80°C.

#### A.3.2 Activity Assay

To measure the kinase activity of these computationally generated ERK sequences, we first set up a kinase reaction with commercial Myline Basic Protein (MBP), a known substrate for ERK1/2 (Seger, 2010). The kinase reaction was performed in kinase buffer (200 mM Tris-HCl pH 8, 100 mM MgCl2, 250 *µ*M DTT, 0.5 mg/ml BSA) 10 *µ*M ATP, 2 *µ*M MBP, and 500 nM of the ERK. We Set the reaction without ERK considered as blank. We also used wild type ERK protein (inactive) purified in our lab as control. The reaction was performed at room temperature for 2 hours.

After the incubation we performed the ADP-Glo Kinase Assay to measure the ADP formed from the kinase reaction (Zegzouti et al., 2009). ADP-Glo is a bioluminescent assay, where it depends on luminescence generation upon ADP conversion to ATP correlating how much light produced to the kinase activity of the protein, as shown in figures S4a, S4b, and S4c. The luminescence of kinase reactions were measured on an infinite M1000Pro plate reader (TECAN). All the reactions were performed in triplicates.

### A.4 Additional ERK2results

**Table 3:** RMSD of the design region (residues 169-188) between designs and the reference structures. Reference structure 4GSB is the target conformation for CSD101, CSD102, MPNN101, and MPNN102 and 2ERK is the target conformation for CSD103, CSD104, MPNN103, and MPNN104. CSD101 and CSD102 show preference for 4GSB over 2ERK, and CSD103 and CSD104 show preference for 2ERK over 4GSB. All ProteinMPNN sequences show preference for 2ERK over 4GSB.

|  | 2ERK | 4GSB |
| --- | --- | --- |
| 2ERK | 0 | 7.102 |
| 4GSB | 7.102 | 0 |
| WT AF3 | 1.787 | 7.392 |
| CSD101 AF3 | 9.593 | 8.151 |
| CSD102 AF3 | 14.196 | 11.092 |
| CSD103 AF3 | 2.046 | 7.549 |
| CSD104 AF3 | 3.115 | 6.831 |
| MPNN101 AF3 | 6.330 | 8.155 |
| MPNN102 AF3 | 6.067 | 7.535 |
| MPNN103 AF3 | 3.535 | 8.297 |
| MPNN104 AF3 | 3.746 | 8.506 |

### A.5 MotifDiv extended results

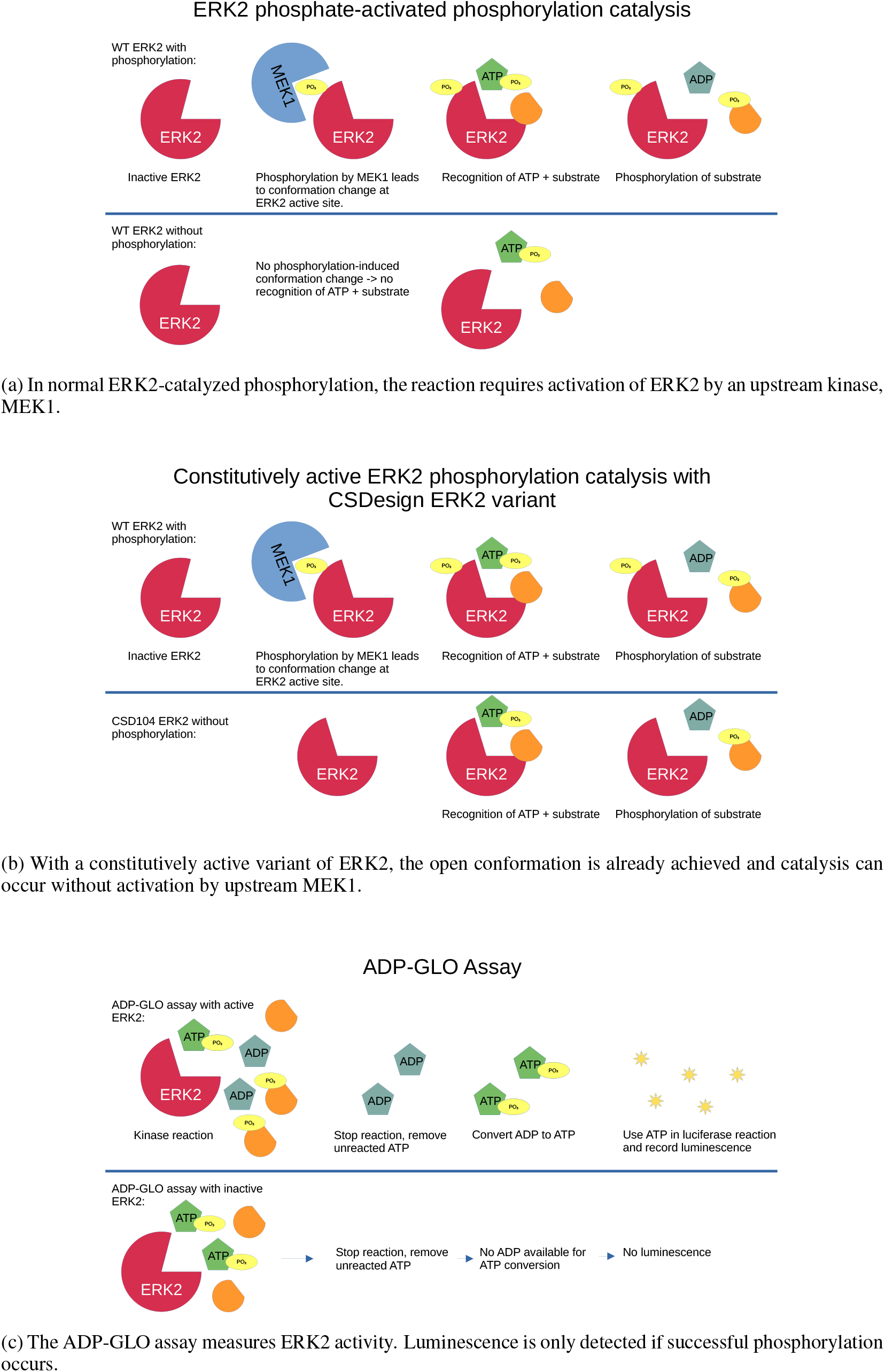

**Figure 5.**
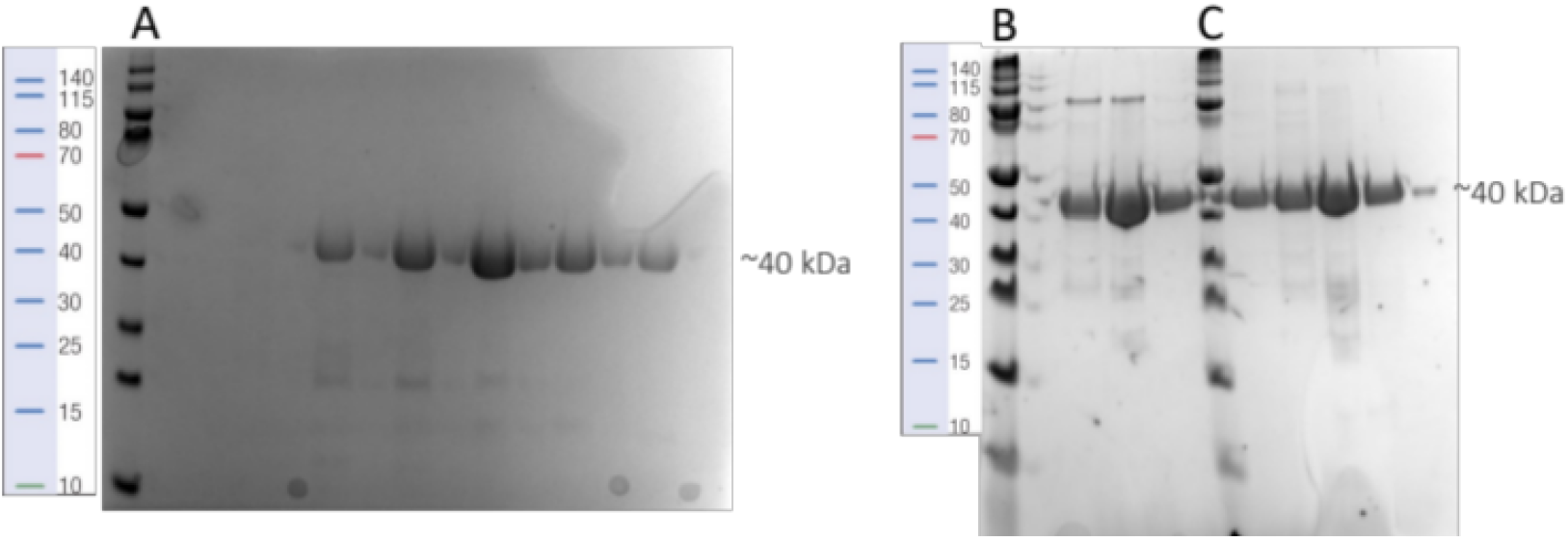
SDS-PAGE representing the fractions collected from the SEC experiments (A) CSD101, (B) CSD103, and (C) CSD104.

**Figure 6.**
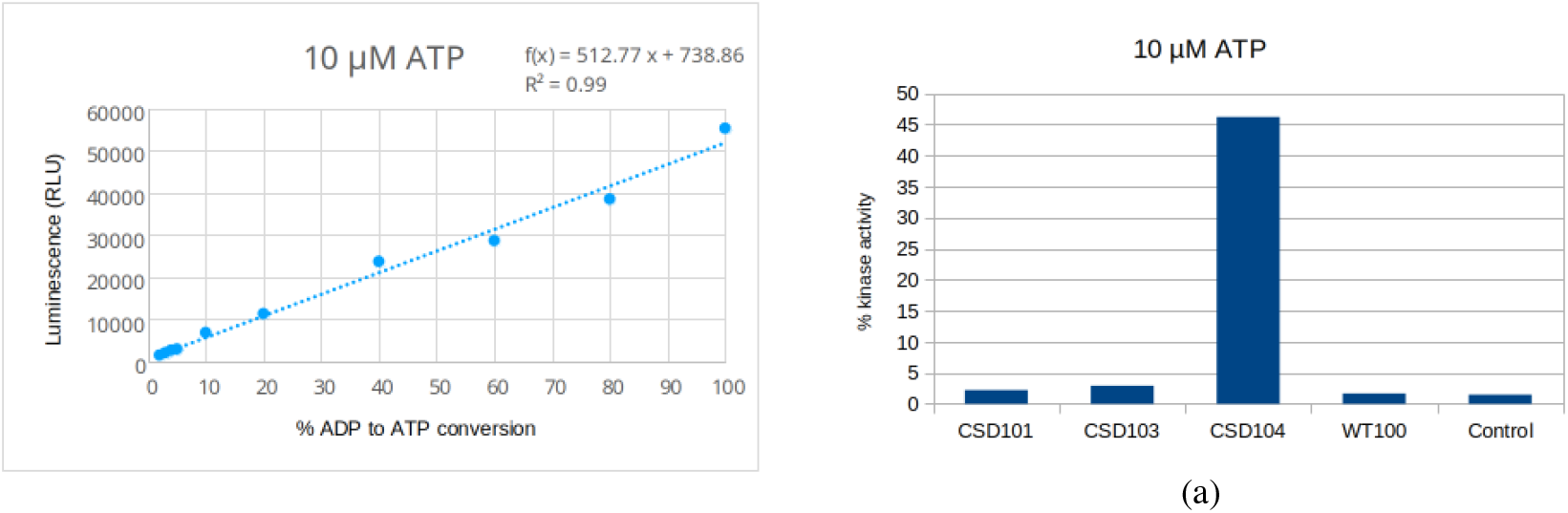
ADP-Glo kinase assay linearity and implementation of the assay for computationally designed ERK constructs. *left:* ADP to ATP standard conversion curve for 10 *µ*M ATP kinase reaction, *right:* ADP-Glo kinase assay for different ERK constructs showing percentage to ATP consumed in the each reaction. All the reactions were performed in triplicates. CSD104 exhibited significant luminescent activity even in the absence of upstream phosphorylation.

**Table 4:**
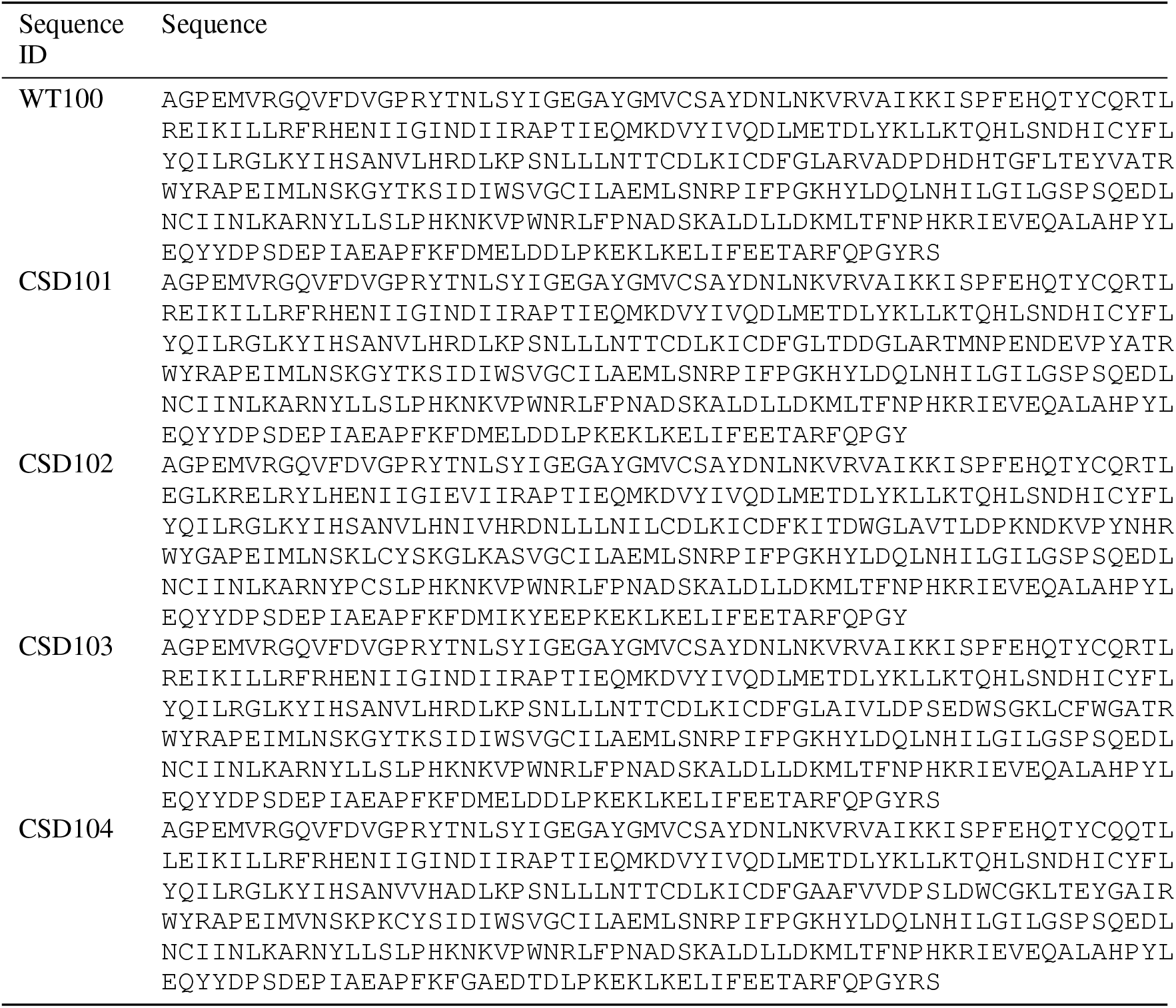
Sequences of proteins included in the ERK2 in vitro assay

**Figure 7.**
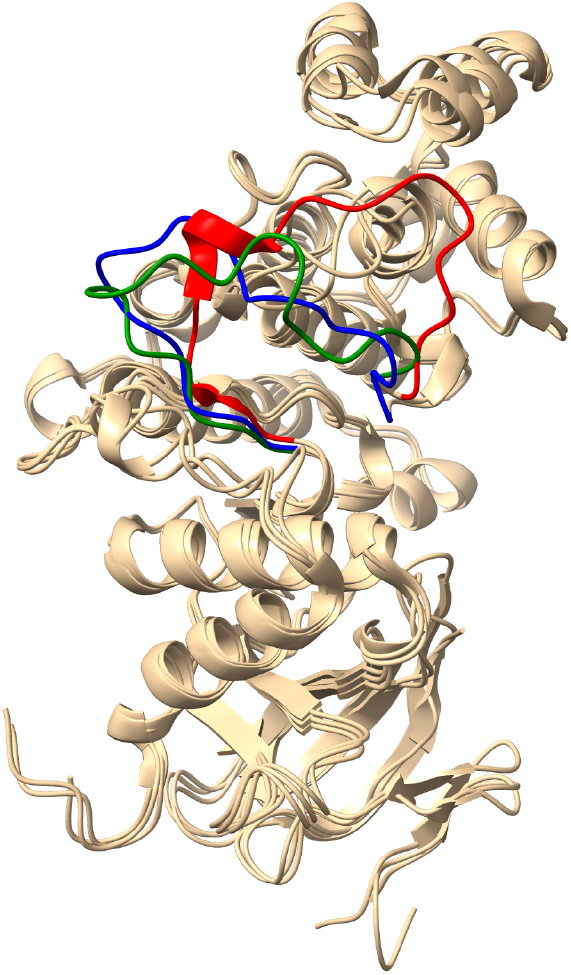
CSD103 (motif shown in blue) showed preference for the active conformation (green) of ERK2 over the inactive conformation (red).

**Figure 8.**
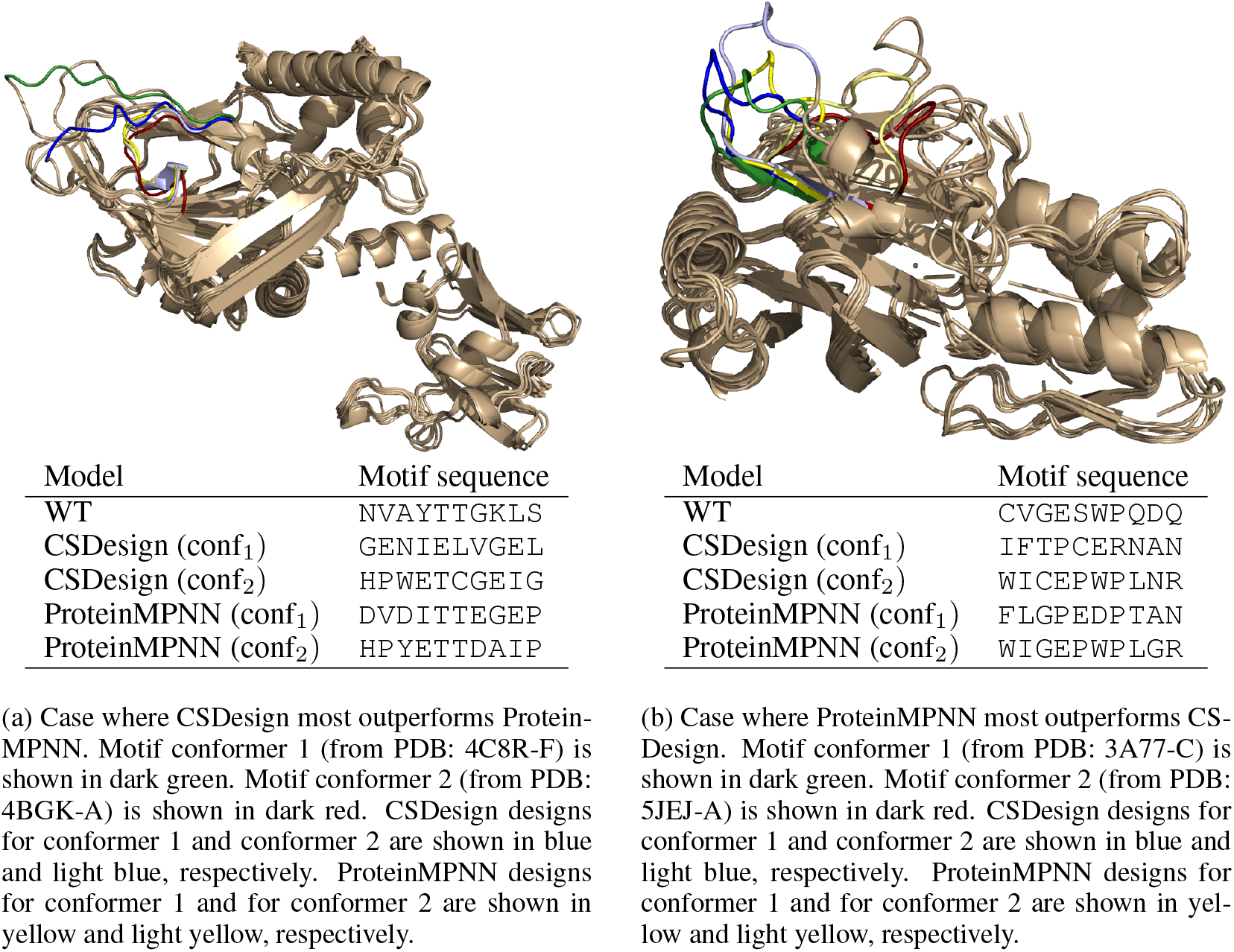
Cases where CSDesign most outperforms ProteinMPNN (left) and Protein-MPNN most outperforms CSDesign (right). “Most outperforms” considers the quantity (*RMSD*_*csd*_(*pred*_1_, *conf*_1_) + *RSMD*_*csd*_(*pred*_2_, *conf*_2_)) − (*RMSD*_*pmpnn*_(*pred*_1_, *conf*_1_) + *RSMD*_*pmpnn*_(*pred*_2_, *conf*_2_)). When this number is positive, it means that CSDesign best recovers both conformers. When this number is negative, it means that ProteinMPNN best recovers both conformers.

## Footnotes

1 This can be seen as a special case of the objective given in Stern et al. (2023), 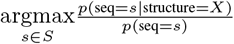 in which *p*(seq = *s*) can be factorized as Σ_*Y*∈*y*_ *p*(seq = *s* structure = *Y*)*p*(structure = *Y*). If *p*(structure = *Z*) = 1, this integral collapses to a single term.

2 This circumvents a limitation of Stern et al. (2023), removing the need for a *p*(seq) model which matches the marginal distribution corresponding to *p*(seq|structure).

## Notes

https://github.com/dellacortelab/cs_design

https://github.com/dellacortelab/motif_div

### Competing Interest Statement

The authors have declared no competing interest.

